# Single-cell transcriptomes of the aging human skin reveal loss of fibroblast priming

**DOI:** 10.1101/633131

**Authors:** Llorenç Solé-Boldo, Günter Raddatz, Sabrina Schütz, Jan-Philipp Mallm, Karsten Rippe, Anke S. Lonsdorf, Manuel Rodríguez-Paredes, Frank Lyko

## Abstract

Fibroblasts are the main dermal cell type and are essential for the architecture and function of human skin. Important differences have been described between fibroblasts localized in distinct dermal layers, and these cells are also known to perform varied functions. However, this phenomenon has not been analyzed comprehensively yet. Here we have used single-cell RNA sequencing to analyze >15,000 cells from a sun-protected area in young and old donors. Our results define four main fibroblast subpopulations that can be spatially localized and functionally distinguished. Importantly, intrinsic aging reduces this fibroblast ‘priming’, generates distinct expression patterns of skin aging-associated genes, and substantially reduces the interactions of dermal fibroblasts with other skin cell types. Our work thus provides comprehensive evidence for a functional specialization of human dermal fibroblasts and suggests that the age-related loss of fibroblast priming contributes to human skin aging.

## Introduction

The skin is the outermost protective barrier of the organism and comprises two main layers, the epidermis and the dermis. The epidermis is a stratified squamous epithelium composed of keratinocytes (around 95%) and other minority cell types such as melanocytes, Merkel and Langerhans cells (Doebel et al., 2017; Shain and Bastian, 2016; Simpson et al., 2011; Woo et al., 2015). The dermis is a much thicker layer located beneath the epidermis and plays an instrumental role in skin architecture and function (Rognoni and Watt, 2018; Sriram et al., 2015). It consists mostly of the extracellular matrix (ECM) generated by its numerous fibroblasts, and also includes many other cell types due to the various structures it harbors, such as the vasculature, nerves, sweat glands and lymphatic vessels (Pawlina and Ross, 2016). A comprehensive molecular characterization of all skin cell types, together with the detailed knowledge of their functions and interactions is crucial to understand skin homeostasis and its role in skin diseases.

Much of our current knowledge about the cellular components of skin has been generated in mice using reporter constructs and lineage tracing, as well as fluorescence-activated cell sorting (FACS) on enzymatically digested tissue. Although limited by the use of predetermined markers, these methods, combined with immunohistochemistry (IHC) have characterized numerous cell (sub)types and defined their location (Alcolea and Jones, 2014; Driskell and Watt, 2015; Kretzschmar and Watt, 2012). These approaches have also described key differences for the fibroblasts in the superficial papillary dermis and the underlying reticular dermis (Driskell and Watt, 2015). For example, while papillary fibroblasts are morphologically thin and spindle-shaped, reticular fibroblasts are squarer and more expanded (Janson et al., 2012; Schafer et al., 1985). Further differences include their rate of proliferation, contractility, production of and response to cytokines and growth factors, as well as the expression of ECM components such as collagens and proteoglycans (Sriram et al., 2015). Examples for the latter include collagen type IV, which is more expressed by papillary fibroblasts, or Decorin and Versican, which are more expressed by papillary and reticular fibroblasts, respectively (Schönherr et al., 1993; Sorrell, 2004). These observations suggest different roles for these two types of fibroblasts and for the different histological layers they define. On the functional level, the papillary dermis (Pawlina and Ross, 2016) is known to be essential for epidermal organization due to its close interactions with keratinocytes (Schumacher et al., 2014; Sriram et al., 2015; Taniguchi et al., 2014). Interestingly, it has been suggested that intrinsic skin aging may have a stronger impact on the papillary fibroblasts, eventually leading to their loss or impaired functionality (Janson et al., 2012; Mine et al., 2008). As a result, the main phenotypic effects observed in old skin could be due to the subsequent impairment of the basal membrane, which plays a key role in epidermal homeostasis (Mine et al., 2008).

Besides their fundamental role in skin architecture, dermal fibroblasts actively participate in cutaneous immune responses, wound healing and communication with the nervous and vascular systems (Haniffa et al., 2007; Sorrell and Caplan, 2009; Werner et al., 2007). This functional diversity is now beginning to be unveiled by single-cell RNA sequencing (scRNA-seq), which allows simultaneous profiling of the transcriptomes of thousands of individual cells. A pioneering study, based on a comparably low number of single, flow-sorted fibroblasts from mouse dorsal skin provided evidence for the heterogeneity of these cells and identified a subtype involved in the fibrotic response to injuries (Rinkevich et al., 2015). Another scRNA-seq study of mid-forearm dorsal whole human skin (2,742 fibroblasts) found two main fibroblast groups displaying different morphology and dermal distribution (*SFRP2*^+^ and *FMO1*^+^), as well as five minor populations (Tabib et al., 2017). Interestingly, one of the main groups (*SFRP2*^+^) included the previously described profibrotic fibroblasts (Rinkevich et al., 2015). However, the analysis was performed with chronically-sun-exposed skin samples from a heterogeneous group of donors. More recently, scRNA-seq was also used to complement a transcriptomic study of bulk flow-sorted and microdissected papillary and reticular fibroblasts from mouse and human dermis (Philippeos et al., 2018). Based on 184 flow-sorted cells from the abdominal skin of a single 64 year-old female donor, the analysis suggested the existence of several fibroblast subgroups (Philippeos et al., 2018). Finally, in another study combining population and single-cell transcriptomics, the analysis of 300 flow-sorted fibroblasts from young and old mice, respectively, detected two cell groups that, upon aging, became less well-defined while presenting lower ECM-related gene expression and a gain of adipogenic traits (Salzer et al., 2018).

We have now analyzed more than 15,000 cells from skin samples that were obtained from a defined, sun-protected area from two "young" (25 and 27 y/o) and three "old" (53-70 y/o) male Caucasian donors. Transcriptomic analysis of the young fibroblasts indicated the presence of four main populations that could be spatially localized and functionally distinguished, consistent with ‘priming’ of fibroblasts into functionally distinct subgroups. Subsequent comparative analysis with the expression profiles obtained from old fibroblasts revealed that all subgroups undergo an age-related loss of their functional and spatial identities. Interestingly, old fibroblast populations also generate specific subsets of skin aging-associated secreted proteins (SAASP) and experience a substantial decrease of their interactions with other skin cell types. Altogether, our work provides a particularly detailed and comprehensive characterization of human dermal fibroblast populations, and provides novel insight into their functional specificities as well as their age-related changes.

## Results

### Single-cell RNA sequencing analysis of sun-protected whole human skin identifies fourteen distinct cell populations

The anatomy of the skin can vary considerably depending on a number of endogenous and environmental factors (Cua et al., 1990; Giacomoni et al., 2009). To minimize confounding effects of these factors, our scRNA-seq analysis was based on five independent whole-skin samples that were obtained specifically from the sun-protected inguinoiliac region of male donors. Since we also sought to address the effects of intrinsic aging on dermal fibroblasts, samples were obtained from two younger (25 and 27 y/o) and three older (53, 69 and 70 y/o) donors. After enzymatically and mechanically disrupting the tissue, dead cells were thoroughly removed and samples were subjected to scRNA-seq using the commercial version of the Drop-seq protocol (Macosko et al., 2015).

In an initial analysis, we obtained an overview of the diverse skin populations by combining the cells from all five samples. Data analysis of the 15,457 cells that passed our quality controls (Table S1, see Methods for details) resulted in a t-distributed Stochastic Neighbor Embedding (t-SNE) plot displaying 14 cell clusters with distinct expression profiles (Figure 1A). Comparing known markers with the most representative expressed genes of each cluster (Figure 1B and Table S2) revealed the identity of the 14 cell clusters, all of which are known constituents of the human skin (Figures 1C and S1). Six clusters comprised the two key cell types of the skin, keratinocytes and fibroblasts. Keratinocytes were detected in two clusters (#6 and #7) and their diversity was mainly due to their degree of differentiation. While epidermal stem cells (EpSC) and other undifferentiated progenitors (#7) expressed markers such as *KRT5*, *KRT14*, *TP63*, *ITGA6* and *ITGB1*, differentiated keratinocytes were defined by *KRT1*, *KRT10*, *SBSN* and *KRTDAP* expression (Moll et al., 2008). Fibroblasts were identified by their archetypal markers *LUM*, *DCN*, *VIM*, *PDGFRA* and *COL1A2* (Philippeos et al., 2018), constituted the most abundant skin cell type (5,913 cells in total), and were represented by four clusters (#1, #2, #3 and #9).

**Figure 1.**
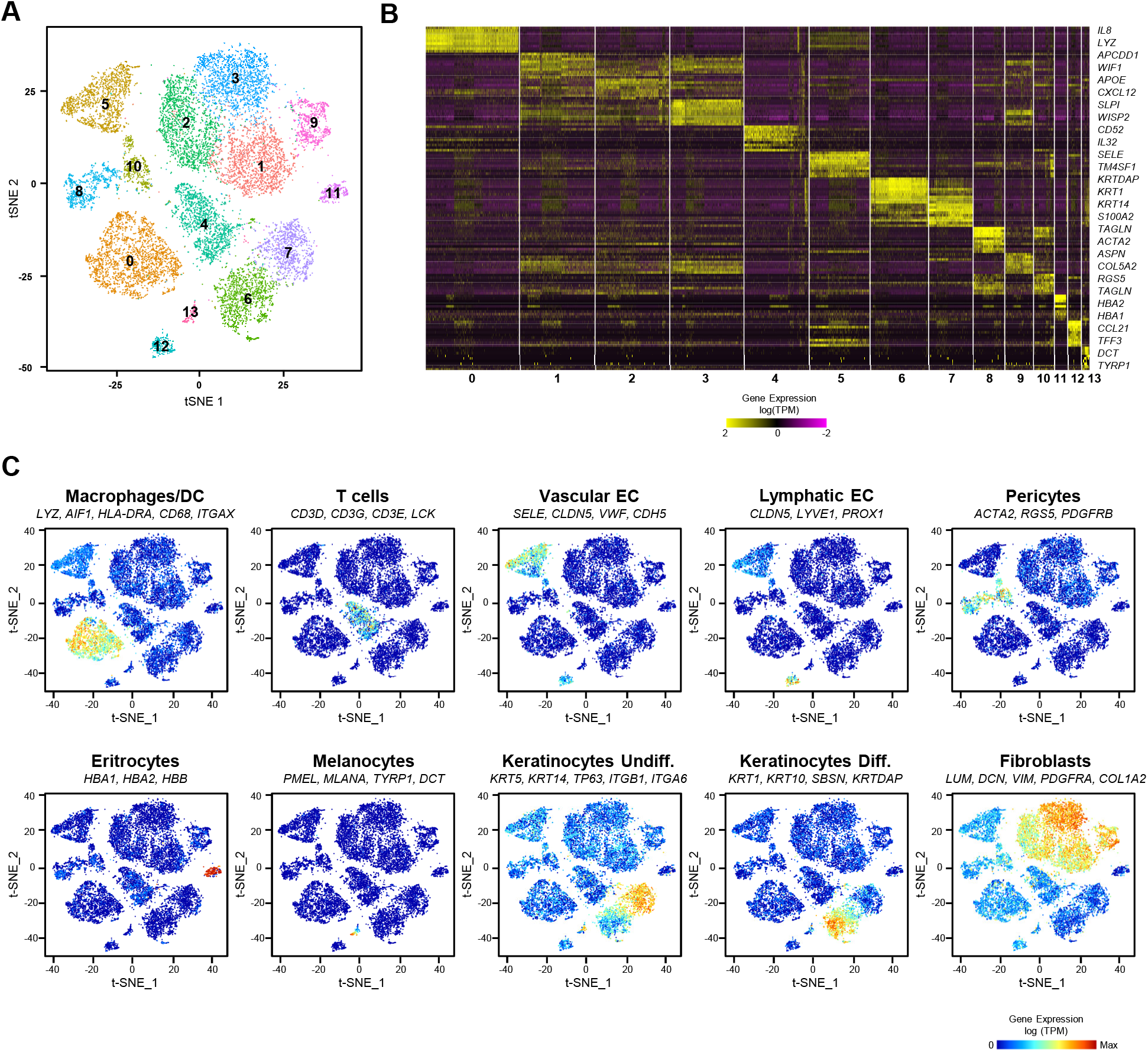
Single-cell RNA sequencing analysis of sun-protected whole human skin identifies fourteen distinct cell populations. **(A)** t-distributed stochastic neighbor embedding (t-SNE) plot depicting single cell transcriptomes from whole human skin (n=5). Each dot represents a single cell (n=15,457). Coloring is according to the unsupervised clustering performed by Seurat. (B) Heatmap showing the ten most differentially expressed genes of each cell cluster, as provided by Seurat. Each column represents a single cell, each row represents an individual gene. Two marker genes per cluster are shown. Yellow indicates maximum relative gene expression and purple indicates no expression. (C) Average expression of 3-5 well-established cell type markers was projected on the t-SNE plot to identify all cell populations (see Methods for details). Red indicates maximum relative gene expression while blue indicates low or no expression. TPM: transcripts per kilobase million.

### Dermal fibroblast populations can be functionally and spatially distinguished

To investigate whether specific functions could be assigned to the different fibroblast subpopulations, we performed gene ontology (GO) analyses using the most representative markers of each cluster. Since it is well established that skin and its fibroblasts undergo specific changes upon aging (Gilchrest, 1996; Rittié and Fisher, 2015; Tigges et al., 2014), we only used the expression profiles of the fibroblasts from young samples (1,795 cells) for this analysis (Figure 2A and Table S3). Classical fibroblast functions related to collagen or ECM production and organization were strongly enriched for three of the clusters (I, III and IV) (Figure 2B). GO analyses also revealed mesenchymal functions such as *skeletal system development, ossification or osteoblast differentiation* for the cells belonging to clusters III or IV (Figure 2B). Interestingly, our results also showed a strong enrichment for functions related to inflammation specifically in the fibroblasts of cluster II. Significant examples include *inflammatory response, cell chemotaxis or negative regulation of cell proliferation*, necessary for the final anchoring of leukocytes (Figure 2B). Functions that are typically attributed to fibroblasts, such as collagen or ECM production and organization, did not appear among the most statistically significant categories for the cells of this cluster. These findings provide a first illustration for the functional heterogeneity of the fibroblasts in our samples.

**Figure 2.**
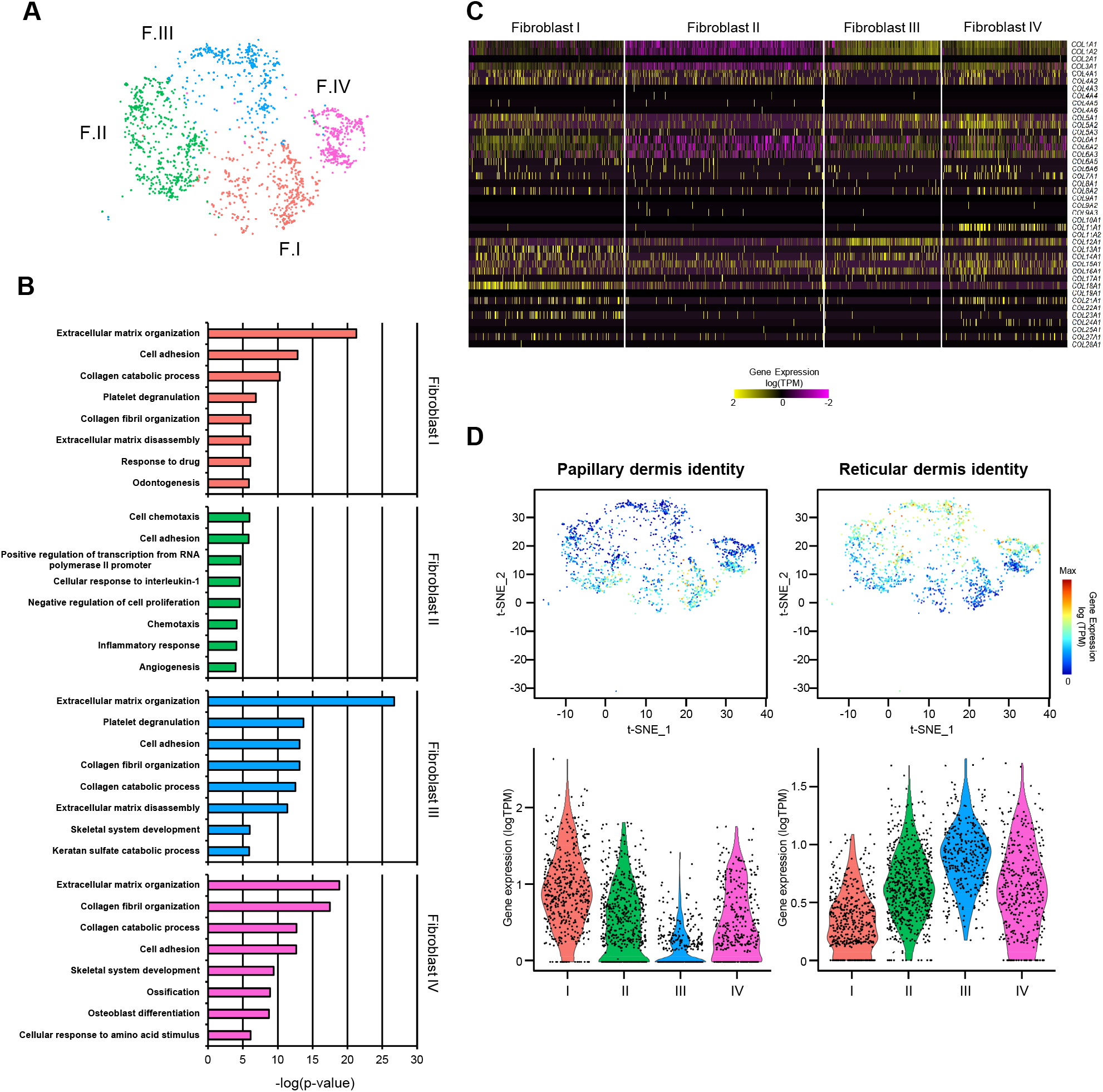
Dermal fibroblast populations can be functionally and spatially distinguished. **(A)** t-SNE plot displaying dermal fibroblasts from young donors (n=2). Each dot represents a single cell (n=1,795). Coloring is according to the unsupervised clustering performed by Seurat. (B) Top 8 enriched Gene Ontology (GO) terms in each fibroblast population sorted by p-value. (C) Heatmap showing the expression of all collagen genes in the distinct fibroblast populations. Each column represents a single cell and each row an individual collagen gene. Yellow indicates maximum relative gene expression while purple indicates no expression. **(D)** Average expression of the genes constituting the papillary and reticular gene signatures for predicting dermal localization of the fibroblasts from the four clusters. In the t-SNE plots, red indicates maximum relative expression and blue indicates low or no expression of a particular set of genes. In the violin plots, X-axes depict cell cluster number and Y-axes represent gene expression in log(TPM). TPM: transcripts per kilobase million.

The expression of some specific collagens has also been linked to particular fibroblast functions (Kadler et al., 2007). We therefore analyzed the four fibroblast clusters at the level of collagen expression patterns. In agreement with our GO analysis, the results clearly indicated lower global collagen expression levels in cluster II (Figure 2C). The analysis also revealed that the collagen genes *COL11A1* and *COL24A1*, associated with cartilage and bone development, respectively (Li et al., 2018; Wang et al., 2012), were specifically expressed in the fibroblasts of cluster IV (Figure 2C), suggesting a stronger mesenchymal component for this cell population. This was also supported by the association of dermal papilla stem cells with this cluster, which represent a well-established population characterized by high *CRABP1* and *TNN* expression levels (Driskell and Watt, 2015; Tabib et al., 2017) (Figure S2). For the remaining collagen and ECM secreting clusters (I and III), the analysis revealed a bias in the production of collagens that have previously been linked to specific dermal locations (Figure 2C). Furthermore, fibroblasts in cluster I express *COL13A1* and *COL23A1*, two known markers of papillary fibroblasts (Haydont et al., 2019; Peltonen et al., 1999; Philippeos et al., 2018; Veit et al., 2011). High expression levels of another epidermal-dermal junction collagen gene, *COL18A1*, also supported the location of these fibroblasts within the papillary layer (Halfter et al., 1998; Haydont et al., 2019; Nauroy et al., 2017; Philippeos et al., 2018).

To better predict the potential localization of the four observed fibroblast populations within the dermis, we next studied the expression of sets of genes that have previously been related to papillary or reticular fibroblasts. While the most representative markers of the papillary fibroblasts comprise *APCDD1*, *AXIN2*, *COLEC12*, *PTGDS* and *COL18A1*, the reticular fibroblast signature is typically defined by a group of ten genes, including *MGP* or *MFAP5* (Haydont et al., 2019; Janson et al., 2012; Nauroy et al., 2017; Philippeos et al., 2018) (Figure S3). In agreement with the collagen expression data, our results showed that the papillary gene expression signature is mostly restricted to fibroblasts in cluster I (Figures 2D and S3). In contrast, fibroblasts in clusters II, III and IV showed more prominent expression of the reticular signature, with the highest reticular expression levels observed in cluster III (Figure 2D).

Taken together, these results reveal ‘priming’ of human skin fibroblasts into four functionally and spatially defined populations. Two populations have prominent roles in the generation of structural collagen and ECM organization. One of these populations is clearly associated with the papillary dermis (cluster I, ‘secretory-papillary fibroblasts’) and the other with the reticular dermis (cluster III, ‘secretory-reticular fibroblasts’). A third population, which is mainly distributed within the reticular dermis, appears to maintain a greater mesenchymal potential and contains the dermal papilla cells (cluster IV, ‘mesenchymal fibroblasts’). Finally, our analysis also identifies a fourth population with pro-inflammatory functions and a mostly reticular localization (cluster II, ‘pro-inflammatory fibroblasts’).

### Immunofluorescence-based detection of fibroblast subpopulations

To further characterize and validate the four fibroblast subpopulations that were identified in our initial analysis, we identified representative markers for each subpopulation according to their expression in specific cell clusters (Table 1). *CCL19*, which encodes the immunoregulatory C-C motif chemokine ligand 19 was selected as a marker for the pro-inflammatory subpopulation (Figure 3A). In addition, *TSPAN8*, which encodes a cell-surface glycoprotein associated with exosomes, was selected as a marker for the secretory-reticular population (Figure 3B). To assess the microanatomical distribution of the characterized subpopulations, we performed immunofluorescence staining of CCL19 and TSPAN8 in healthy human skin sections together with Vimentin, a general marker for all fibroblast populations. The results showed that the CCL19^+^/VIM^+^ pro-inflammatory cells are present in both the papillary and reticular dermis (Figure 3C). Also, the detection of TSPAN8^+^/VIM^+^ cells confirmed the presence of the secretory-reticular population of fibroblasts in the reticular dermal region (Figure 3D). These results provide important confirmation for our findings obtained by single-cell transcriptomics and identify first markers for the immunohistochemical analysis of specific fibroblast subpopulations.

**Table 1.** Representative marker genes of each fibroblast populations. The table depicts the top 5 genes selected as marker genes of each fibroblast population according to their fold-change and enriched expression in a particular population.

|  | Gene | Fold-change | % cells in cluster | % cells not in cluster | Adjusted p-value |
| --- | --- | --- | --- | --- | --- |
| **Secretory**<br>**Papillary** | APCDD1 | 4.66 | 0.768 | 0.167 | 3.44E-139 |
|  | WIF1 | 3.40 | 0.49 | 0.079 | 2.73E-85 |
|  | ID1 | 3.07 | 0.616 | 0.238 | 5.62E-59 |
|  | COL18A1 | 2.95 | 0.591 | 0.172 | 3.44E-74 |
|  | NKD2 | 2.29 | 0.384 | 0.064 | 1.49E-60 |
| **Proinflammatory** | CCL19 | 15.73 | 0.338 | 0.077 | 3.39E-47 |
|  | CXCL3 | 4.70 | 0.528 | 0.285 | 7.55E-30 |
|  | CYGB | 2.24 | 0.374 | 0.062 | 1.02E-59 |
|  | RARRES2 | 2.06 | 0.672 | 0.265 | 2.89E-59 |
|  | GGT5 | 1.96 | 0.317 | 0.069 | 1.35E-40 |
| **Secretory**<br>**Reticular** | WISP2 | 13.47 | 0.877 | 0.132 | 2.24E-207 |
|  | SLPI2 | 10.14 | 0.764 | 0.095 | 4.67E-173 |
|  | CTHRC1 | 7.54 | 0.821 | 0.111 | 2.18E-187 |
|  | PCSK1N | 5.25 | 0.541 | 0.062 | 1.06E-111 |
|  | TSPAN8 | 3.99 | 0.624 | 0.107 | 3.44E-107 |
| **Mesenchymal** | ASPN | 6.99 | 0.646 | 0.118 | 1.42E-113 |
|  | POSTN | 4.39 | 0.619 | 0.131 | 1.02E-87 |
|  | TNN | 3.37 | 0.339 | 0.001 | 1.86E-107 |
|  | SFRP1 | 2.56 | 0.389 | 0.09 | 2.67E-45 |
|  | GPC3 | 2.43 | 0.5 | 0.169 | 3.63E-40 |

**Figure 3.**
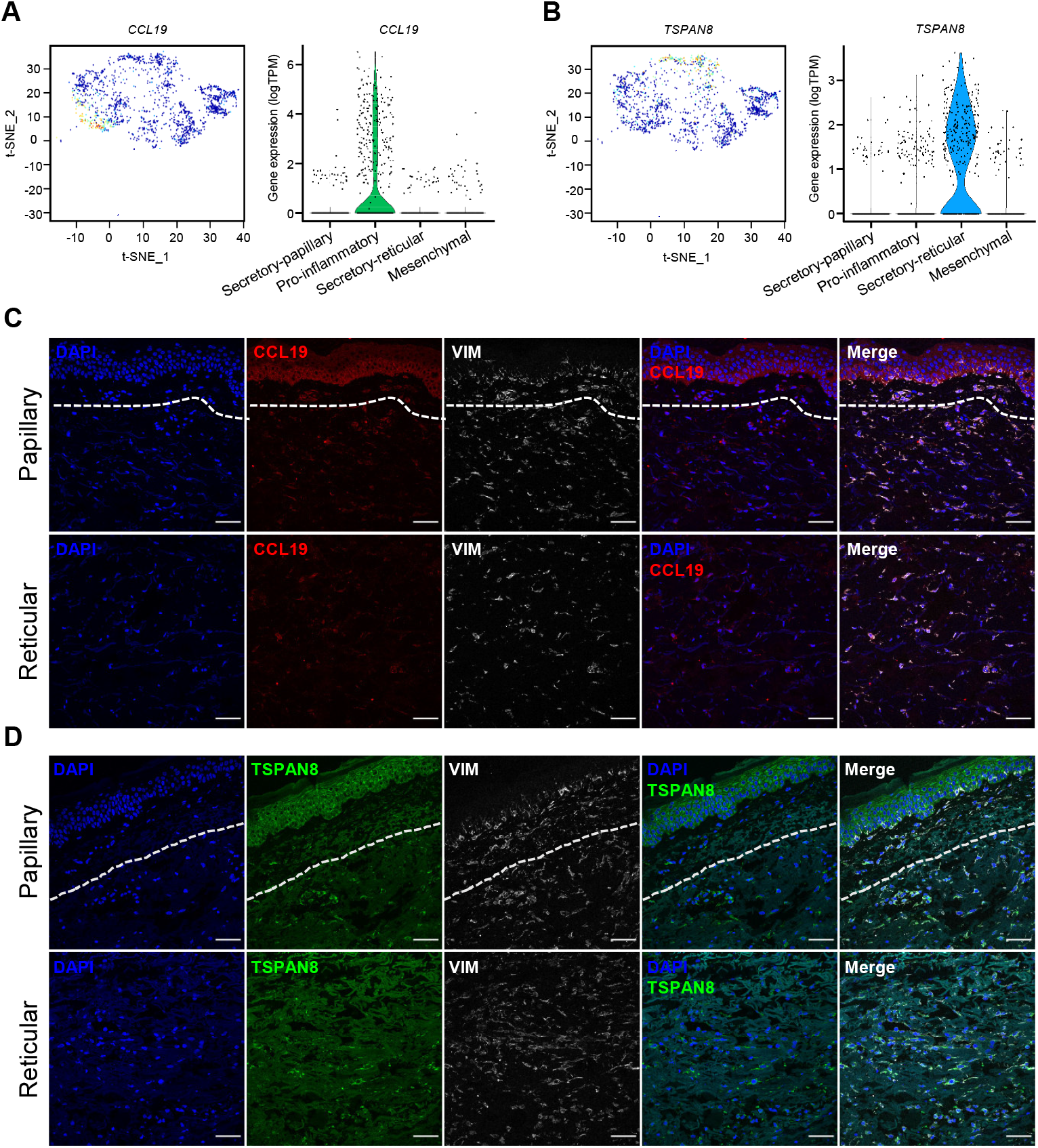
Immunofluorescence stainings detect the pro-inflammatory and secretory-reticular fibroblast subpopulations. t-SNE and violin plots showing population-specific expression levels for the secretory-reticular and pro-inflammatory markers CCL19 (A) and TSPAN8 (B), respectively. In the t-SNE plots, red indicates maximum relative expression and blue indicates low or no expression of a particular gene. In the violin plots, the X-axis depicts cell cluster identity while the Y-axis represents gene expression in log(TPM). (C,D) Representative confocal images of human skin sections stained for CCL19 (red, C) TSPAN8 (green, D) and VIM (grey). Papillary and reticular (superficial and deep) dermal regions are indicated. Nuclei were stained with DAPI (blue). Dotted lines denote boundaries between papillary and reticular dermis. Images are shown at 40× original magnification. Scale bar, 50 μm. TPM: transcripts per kilobase million.

### Aging leads to a loss of dermal fibroblast priming

We also investigated the effect(s) of aging at the level of dermal fibroblast populations. To this end, we stratified each population into old and young fibroblasts (Figure S4). Consistent with the reduced proliferative capacity of aged cells, old fibroblasts from all four subpopulations showed a delay at the G1/S transition of the cell cycle (Figure 4A). Subsequent GO analyses of the most representative genes defining the old populations (Table S4) revealed a considerable age-dependent loss of functional identity for each cluster. In comparison to young fibroblast subpopulations, the aged counterparts showed fewer function-related terms, and substantially reduced p-values, consistent with fewer genes supporting the terms (Figure 4B). A similar effect was also seen at the level of collagen production, which became particularly decreased in the secretory clusters (Figures 4C and S5). Finally, aging also affected spatial gene expression signatures, as papillary fibroblasts became less papillary and more reticular, while reticular fibroblasts presented a less prominent reticular signature (Fig. 4D). Taken together, these findings are in agreement with an age-related loss of fibroblast priming.

**Figure 4.**
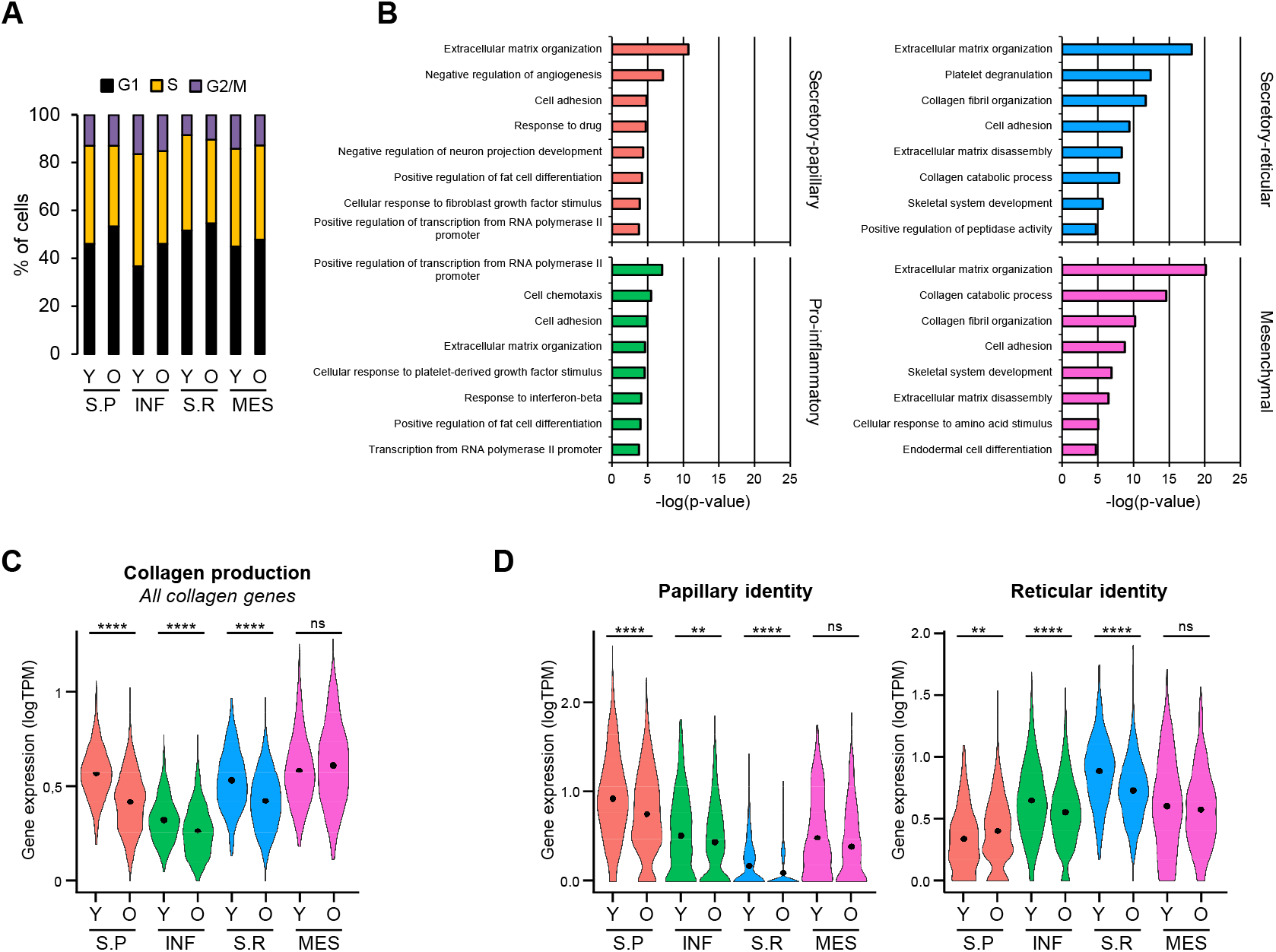
Aging leads to loss of dermal fibroblast priming. **(A)** Percentage of fibroblasts of each population that were in the G1, S or G2/M phase of the cell cycle in young and old skin samples, respectively. (B) Top 8 enriched Gene Ontology (GO) terms in each old fibroblast population sorted by p-value. Coloring is according to the unsupervised clustering performed by Seurat. (C) Violin plots displaying the average expression of all collagen genes in the fibroblasts of all populations, for young and old skin. (D) Violin plots displaying the expression of the papillary and reticular identity gene signatures in the fibroblasts of all populations, for young and old skin. In violin plots, X-axes depict cell cluster number and Y-axes represent gene expression in log(TPM). Cluster average expression of each set of genes is shown as a black dot. Statistical analysis was performed using a Wilcoxon Rank Sum test (*p < 0.05, **p < 0.01, ***p < 0.001, ****p < 0.0001). TPM: transcripts per kilobase million; Y: young; O: old; S.P: Secretory-Papillary; INF: Pro-inflammatory; S.R: Secretory-Reticular; MES: Mesenchymal.

### Old fibroblasts express population-specific SAASP-associated genes and show a disrupted cell-cell interactome

Finally, we also analyzed whether age-related changes in fibroblast populations could explain age-related skin phenotypes. For example, it is known that aged fibroblasts become more susceptible to the accumulation of reactive oxygen species (Kozieł et al., 2011). Interestingly, our GO analyses performed with the most down-regulated genes of each aged cluster show that genes related to hydrogen peroxide metabolism are decreased in all fibroblast populations (Figure S6). Furthermore, old skin is known to acquire a chronic, low-grade inflammatory phenotype (Zhang and Duan, 2018; Zhuang and Lyga, 2014). This could also be detected at the level of fibroblast populations, as we find *immune response* among the main enriched terms in our GO analyses of the most up-regulated genes of each aged cluster (Figures S6 and S7). Old fibroblasts also showed changes in the expression of genes that were previously detected in the aging dermal fibroblast secretome (Waldera Lupa et al., 2015). Furthermore, differential expression profiles of SAASPs could also be observed in the different old fibroblast populations (Figures 5A and S8). Taken together, our scRNA-seq data thus recapitulates known phenotypes associated with skin aging.

**Figure 5.**
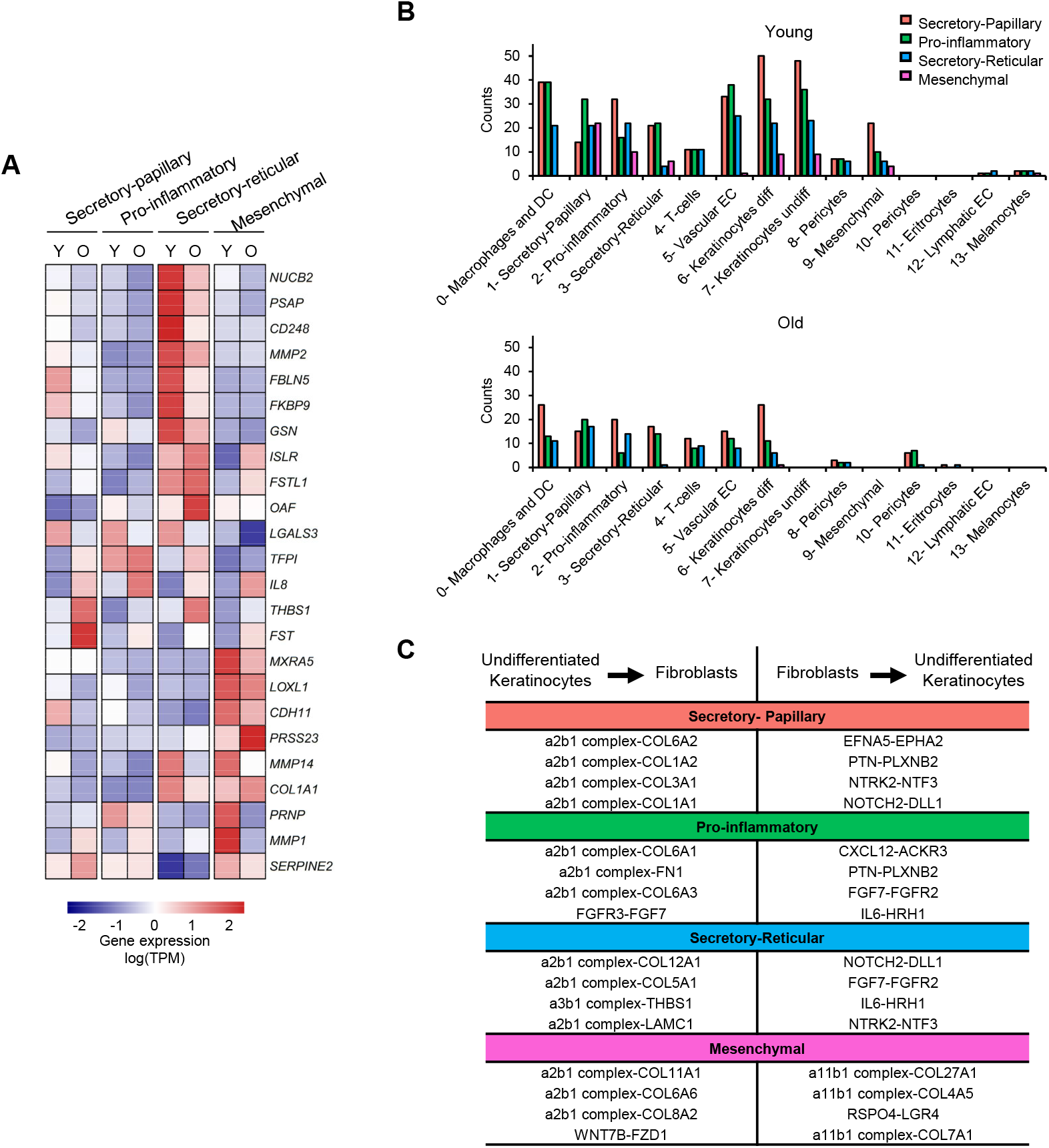
Old fibroblasts express population-specific SAASP-associated genes and show a disrupted cell-cell interactome. **(A)** Expression of SAASP genes (rows) that are differentially (fc >1.25) expressed between young and old fibroblasts in at least one population (columns). The heatmap shows the mean relative expression, scaled by row (gene). (B) Bar plots showing the number of interactions established by the four observed fibroblast populations with the rest of the cell types identified in human skin, in young (up) and old (down) samples. Coloring and numbering is according to the original unsupervised clustering performed by Seurat. (C) Summary of the top 4 interactions lost between each fibroblast population and undifferentiated keratinocytes, sorted by *p-value*. The table depicts interactions in both directions for each pair. TPM: transcripts per kilobase million. Y: young; O: old; S.P: Secretory-Papillary; INF: Pro-inflammatory; S.R: Secretory-Reticular; MES: Mesenchymal.

Finally, fibroblasts are known to establish interactions with many other skin cell types during homeostasis (Sriram et al., 2015). Importantly, scRNA-seq also provides novel opportunities to identify communicating pairs of cells based on the expression of cell-surface receptors and their interacting ligands (Vento-Tormo et al., 2018). Interestingly, our results indicate that most interactions identified in young fibroblasts are potentially lost during intrinsic skin aging. This effect was particularly pronounced for interactions involving undifferentiated keratinocytes (Figure 5B and specific examples are shown in Figure 5C) These findings suggest that the loss of interactions between fibroblasts and communicating cell types represent a major and previously unrecognized molecular phenotype of the aging skin.

## Discussion

Single-cell transcriptomics currently represents the most effective method to define cell populations in a given tissue (Deng et al., 2019, Kolodziejczyk et al., 2015.). However, whole human skin has not been analyzed at a comprehensive level yet, as previous single-cell studies were either focused on sun-exposed material obtained from a heterogeneous group of donors (Tabib et al., 2017) or provided limited coverage from flow-sorted cells (Philippeos et al., 2018). Our study analyzes single-cell transcriptomes from more than 15,000 skin cells that were all obtained from the same sun-protected location and only from healthy male donors. This allowed us to minimize confounding effects and provide the first description of age-related changes in human dermal fibroblasts.

We describe an overall cellular composition of the human skin samples that is consistent with previous observations (Tabib et al., 2017). In addition, we also describe several fibroblast subpopulations with distinct functional and spatial specifics. These findings provide an important extension of previous observations describing fibroblast heterogeneity, including the distinction of papillary and reticular fibroblasts (Driskell and Watt, 2015; Lynch and Watt, 2018; Sriram et al., 2015). More specifically, our results suggest the existence of four major dermal fibroblast populations: secretory-papillary fibroblasts, secretory-reticular fibroblasts, mesenchymal fibroblasts and pro-inflammatory fibroblasts.

Interestingly, the attributes of the four populations all reflect known functions of fibroblasts. For example, the secretion of collagens and extracellular matrix components is considered as the defining function of dermal fibroblasts, with well-known differences in the secretory activities of papillary and reticular fibroblasts. In agreement, our results define secretory-papillary and secretory-reticular fibroblasts as two separate subpopulations. Similarly, it is well known that fibroblasts can differentiate into other mesenchymal cell types (Chen et al., 2007; Kuroda et al., 2010), which is reflected by our mesenchymal population. We notice that we did not detect any evidence for adipogenic differentiation in our dataset, which is likely related to the specific characteristic of the sampling site (inguinoiliac region of male donors). Finally, the pro-inflammatory functions of dermal fibroblasts are also well established (Kendall and Feghali-Bostwick, 2014) and correspondingly reflected in our pro-inflammatory fibroblast subpopulation. Our results thus suggest that the functional heterogeneity of fibroblasts can be explained by the existence of functionally primed subpopulations.

Our secretory-papillary and secretory-reticular fibroblasts are defined by the expression of previously established markers for these two dermal layers. The pro-inflammatory fibroblasts presented a mixed signature, which was also supported by the widespread localization of the *CCL19*^+^ pro-inflammatory fibroblasts. These findings are in agreement with the notion that the entire dermis may require the protective function of pro-inflammatory fibroblasts. Similarly, the mesenchymal fibroblasts also displayed a mixed localization signature, which may indicate different localizations of specific subpopulations. For example, dermal papilla stem cells expressed a papillary dermis signature (Figure S9), consistent with their shared origin with papillary fibroblasts (Driskell and Watt, 2015).

We notice that some of our findings show differences to previous studies. For example, Tabib et al. described only two major fibroblast populations that were defined by the expression of *SFRP2* and *FMO1*, as well as other five minor, closely related populations (Tabib et al., 2017). However, these populations were not functionally defined and their significance remained unclear. In our dataset, expression of *FMO1* was very low in fibroblasts, while *SFRP2* was expressed by both secretory populations and a subgroup of the pro-inflammatory population (Figure S10). While the reasons for these discrepancies remain to be elucidated, it is possible that the lower number of cells, in combination with sampling from a different, sun-exposed region (dorsal forearm) may have resulted in a less accurate stratification of fibroblast subpopulations. On the other hand, the scRNA-seq experiment performed by *Philippeos et al*. with only 184 flow-sorted fibroblasts obtained from a single abdominal skin sample detected five subpopulations (Philippeos et al., 2018). While the two major subpopulations expressed markers that might localize them in different dermal layers, the significance of the minor subpopulations remained unclear. One of these subpopulations comprised only 5 cells, two others appeared to contain pericytes and pre-adipocytes, respectively (Philippeos et al., 2018).

Our results also suggest a pronounced age-related loss of fibroblast priming. This was detectable both at the level of genes defining their functions, and in the expression of their spatial localization signatures. These findings provide an important complement to a recent study that described age-related identity loss in murine fibroblasts (Salzer et al., 2018). While we also observed an upregulation of genes related to immune response and inflammation, we did not detect an upregulation of adipogenesis genes. These similarities and differences are likely explained by the limited evolutionary conservation of mouse and human skin (Wong et al., 2011).

Fibroblasts maintain various paracrine interactions with other skin cell types, as well as direct cell-cell interactions (Lynch and Watt, 2018; Sriram et al., 2015). For instance, their contacts along the dermal-epidermal junction with the epidermal stem and progenitor cells (EpSPCs) are key for proper epidermal homeostasis (Schumacher et al., 2014; Sriram et al., 2015; Taniguchi et al., 2014). Importantly, our analysis of the interactome of each fibroblast subpopulation shows that aging causes a considerable decrease in all their potential interactions, and especially those that are maintained with undifferentiated keratinocytes. This may provide a plausible explanation for the thinning and flattening of the dermal-epidermal junction, which constitutes one of the most important phenotypic effects of aged skin (Mine et al., 2008).

Taken together, our study reveals an important pattern of fibroblast heterogeneity at the level of cellular subpopulations and provides a novel concept to understand the role of fibroblasts in skin aging.

## Methods

### Clinical samples

Skin specimens were obtained from patients undergoing routine surgery at the Department of Dermatology, University Hospital of Heidelberg. Only remnant, clinically healthy skin, not required for diagnostic purposes, was analyzed after written informed consent by the patient and as approved by the Ethics Committee of Heidelberg University (S-091/2011) in compliance with the current legislation and institutional guidelines.

### Single-cell RNA sequencing

For each experiment, 4-mm punch biopsies were obtained from healthy whole skin specimens, immediately after resection from the inguinoiliac region of five male subjects. Donor’s characteristics are summarized in Table S1. Samples were kept in MACS Tissue Storage Solution (Miltenyi Biotec) for no longer than 1 h before their enzymatical and mechanical dissociation with the Whole Skin Dissociation kit for human material (Miltenyi Biotec) and the Gentle MACS dissociator (Miltenyi Biotec), following the manufacturer’s instructions. Cell suspensions were then filtered through 70-µm cell strainers (Falcon) and depleted of apoptotic and dead cells with the Dead Cell Removal Kit (Miltenyi Biotec).

Sequencing libraries were subsequently prepared following the Drop-seq methodology (Macosko et al., 2015), using a Chromium Single Cell Controller and the v2 chemistry from 10X Genomics. Thus, approximately 20,000 cells per sample were mixed with the retrotranscription reagents and pipetted into a Chip A Single Cell, also containing the Single Cell 3’ Gel Bead suspension and Partitioning Oil. The Chip was subsequently loaded into a Chromium Single Cell Controller (10X Genomics) where the cells were captured in nanoscale droplets containing both the reagents needed for reverse transcription and a gel bead. Resulting gel bead-in-emulsions (GEMs) were then transferred to a thermocycler in order to perform the retrotranscription following the manufacturer’s protocol. Each gel bead contained a specific 10X Genomics barcode, an Illumina R1 sequence, a Unique Molecular Identifier (UMI) and a poly-dT primer sequence. Therefore, from poly-adenylated mRNA the reaction produced full-length cDNA with a unique barcode per cell and transcript, which allowed tracing back all cDNA coming from each individual cell. Following an amplification step, cDNA was further processed by fragmentation, end repair and A-tailing double-sided size selection using AMPure XP beads. Finally, Illumina adaptors and a sample index were added through PCR using a total number of cycles adjusted to the cDNA concentration. After sample indexing, libraries were again subjected to double-sided size selection. Quantification of the libraries was carried out using the Qubit dsDNA HS Assay Kit (Life Technologies), and cDNA integrity was assessed using D1000 ScreenTapes (Agilent Technologies). Paired-end (26+74bp) sequencing (100 cycles) was finally performed with a HiSeq 4000 device (Illumina).

### Data analysis

Raw sequencing data was processed with Cell Ranger, version 2.1.0, from 10X Genomics. For downstream analysis of the data we used the Seurat package version 2.3.4 (Satija et al., 2015) in R version 3.5.1 (R Core Team, 2018). 16,062 cells passed the quality control steps performed by Cell Ranger. For further quality control, we filtered out cells having less than 10 or more than 25,000 genes expressed and more than 5% mitochondrial reads. The application of this filters resulted in a final dataset of 15,457 single cell transcriptomes.

We stratified our samples into young (n=2) and old (n=3) according to their age at the moment of excision. Both datasets were initially treated separately for log normalization and scaling of the data using Seurat functions NormalizeData() and ScaleData(). Afterwards, the data of both age groups was merged and dimensionality was reduced using a canonical correlation analysis (CCA) implemented in the Seurat package. For subsequent analyses, we selected 20 correlation components (CCs) as they explained more variability than expected by chance using a shared correlation strength approach. We used these aligned CCs as input for clustering with the graph-based method implemented in Seurat and for t-distributed stochastic neighbor embedding (t-SNE) visualization. Using the standard resolution value of 0.6 we identified 14 distinct cell clusters in our complete dataset.

To identify the genes whose expression is enriched in each cell cluster we used the FindAllMarkers() function in each age group separately. This function uses a Wilcoxon Rank Sum test to identify the representative genes of each cluster. These representative genes were used to establish the cell identity of each cluster together with markers found in the literature for cell types typically present in the human skin. Mean expression of a particular set of marker genes per cell was used for cell type identification and was projected into t-SNE or Violin plots.

Gene expression signatures used for the definition of cell populations were: *ACTA2*, *RGS5* and *PDGFRB* (pericytes, clusters #8 and #10) (Paquet-Fifield et al., 2009), *KRT5*, *KRT14*, *TP63*, *ITGB1* and *ITGA6* (epidermal stem cells and other undifferentiated progenitors, cluster #7), *KRT1*, *KRT10*, *SBSN* and *KRTDAP* (differentiated keratinocytes, cluster #6), *PDGFRA*, *LUM*, *DCN*, *VIM* and *COL1A2* (fibroblasts, clusters #1, #2, #3 and #9), *AIF1*, *LYZ*, *HLA-DRA*, *CD68* and *ITGAX* (macrophages and dendritic cells (DC), cluster #0) (Elizondo et al., 2017), *CD3D*, *CD3G*, *CD3E* and *LCK* (T cells, cluster #4) (Chtanova et al., 2005), *SELE*, *CLDN5*, *VWF* and *CDH5* (vascular endothelial cells, cluster #5) (Leeuwenberg et al., 1992), *PROX1*, *CLDN5* and *LYVE1* (lymphatic endothelial cells, cluster #12) (Lee et al., 2015), *HBA1*, *HBA2* and *HBB* (erythrocytes, cluster #11) (Cohen et al., 2008) and *PMEL*, *MLANA*, *TYRP1* and *DCT* (melanocytes, cluster #13) (Bissig et al., 2016).

In order to analyze the genes differentially expressed by fibroblast clusters upon aging we used the FindMarkers() function, comparing the young and old cells present in each cluster.

For the gene ontology (GO) analyses, representative genes expressed by each fibroblast cluster or genes differentially expressed in each cluster upon aging were queried into the Gene Functional Annotation Tool from the DAVID Bioinformatics Database (version 6.8) in order to link each list of genes to relevant biological processes. Gene ontology option GOTERM_BP_ALL was selected and the first GO 8 terms with a p-value <0.05 were chosen as significant categories.

Average gene expression for collagen genes and for a set of genes defining spatial identities was calculated in young and old fibroblast populations and projected onto t-SNE or Violin plots. To test for significance, a Wilcoxon Rank Sum test was used.

To analyze the putative cell-cell interactions established by the distinct cell types present in the human skin we used the publicly available repository CellPhoneDB (Vento-Tormo et al., 2018). This approach performed pairwise comparisons between all the clusters present in our dataset. Only receptors or ligands expressed by at least 10% of the cells in a cluster were used in the analysis. Interactions with a p-value < 0.05 were selected as significant.

### Immunohistochemistry

Remnant, healthy whole skin specimens were fixed overnight in 4% formalin in PBS, paraffin-embedded and cut into 4 µm sections. Then, sections were deparaffinized in xylene and rehydrated in a gradient of ethanol and distilled water prior to heat-induced antigen retrieval. Slides were incubated for 30 minutes at 95 ºC in a water bath in 10 mM Citrate buffer (pH 6.0) containing 0.05% Tween-20.

Skin sections were permeabilized by incubation with 0.4% Triton-X in 1 % Normal Goat Serum (NGS) for 10 minutes, twice. Subsequently, non-specific antibody binding was blocked by incubation with 10% NGS for 1 h followed by overnight incubation with primary antibodies diluted in blocking solution at 4 ºC. Primary antibodies used were rabbit anti-Tspan8 (Abcam, Ab230448, 1:200), mouse anti-Ccl19 (R&D Systems, MAB361, 15 µg/ml), rabbit anti-Vimentin (Cell Signaling, D21H3, 1:100) and chicken anti-Vimentin (Abcam, Ab24525, 1:2000). After washing with PBS-0.1% Tween-20, a second blocking step was performed with 10 % NGS for 10 minutes. Sections were then incubated with corresponding Alexa Fluor-conjugated secondary antibodies (Life Technologies) for 2 h at room temperature. Nuclear counterstaining was performed with DAPI and slides were mounted with Vectashield (Vector Laboratories). Images were taken with a Leica TCS SP5 (Leica Microsystems) confocal microscope using a 40X oil immersion lens and further processed using the Fiji software (Schindelin et al., 2012).

### Data access

scRNA-seq datasets are available from the Gene Expression Omnibus (GEO) database and a reviewer access will be provided upon request.

## Acknowledgements

We thank Dirk Tönges and Miguel Sánchez Eimil for their dedicated support and Katharina Röck and Marc Winnefeld for helpful advice and stimulating discussions. We also thank Felix Bormann for computational support. This work was supported by a research grant from the Helmholtz program ‘Aging and Metabolic Programming’ (AMPro) to F.L.

## Author Contributions

L.S.-B., G.R., A.S.L., M.R.-P. and F.L. analyzed the data. L.S.-B. performed scRNA-seq with support from J.-P.M. and K.R. S.S and L.S.-B. performed IHC assays. A.S.L. provided clinical samples. M.R.-P. and F.L. conceived the study. L.S.-B., M.R.-P. and F.L. wrote the paper with input from other authors. All authors read and approved the final manuscript.

## References

Alcolea, M.P., and Jones, P.H. (2014). Lineage analysis of epidermal stem cells. Cold Spring Harb. Perspect. Med. 4, a015206.

Bissig, C., Rochin, L., and van Niel, G. (2016). PMEL amyloid fibril formation: the bright steps of pigmentation. Int. J. Mol. Sci. 17, pii: E1438.

Chen, F.G., Zhang, W.J., Bi, D., Liu, W., Wei, X., Chen, F.F., Zhu, L., Cui, L., and Cao, Y. (2007). Clonal analysis of nestin (-) vimentin (+) multipotent fibroblasts isolated from human dermis. J. Cell Sci. 120, 2875–2883.

Chtanova, T., Newton, R., Liu, S.M., Weininger, L., Young, T.R., Silva, D.G., Bertoni, F., Rinaldi, A., Chappaz, S., Sallusto, F., et al. (2005). Identification of T cell-restricted genes, and signatures for different T cell responses, using a comprehensive collection of microarray datasets. J. Immunol. 175, 7837–7847.

Cohen, R.M., Franco, R.S., Khera, P.K., Smith, E.P., Lindsell, C.J., Ciraolo, P.J., Palascak, M.B., and Clinton, H.J. (2008). Red cell life span heterogeneity in hematologically normal people is sufficient to alter HbA1c. Blood 112, 4284–4291.

Cua, A., Wilhelm, K., and Maibach, H. (1990). Elastic properties of human skin: relation to age, sex and anatomical region. Arch. Dermatol. Res. 282, 283–288.

Deng Y., Bao F., Dai Q., Wu L.F., and Altschuler S.J (2019). Scalable analysis of cell-type composition from single-cell transcriptomics using deep recurrent learning. Nat. Methods 16, 311–314.

Doebel, T., Voisin, B., and Nagao, K. (2017). Langerhans Cells – The Macrophage in Dendritic Cell Clothing. Trends Immunol. 38, 817–828.

Driskell, R.R., and Watt, F.M. (2015). Understanding fibroblast heterogeneity in the skin. Trends Cell Biol. 25, 92–99.

Elizondo, D.M., Andargie, T.E., Yang, D., Kacsinta, A.D., and Lipscomb, M.W. (2017). Inhibition of allograft inflammatory factor-1 in dendritic cells restrains CD4^+^ T cell effector responses and induces CD25^+^ Foxp3^+^ T regulatory subsets. Front. Immunol. 8, 1502.

Giacomoni, P.U., Mammone, T., and Teri, M. (2009). Gender-linked differences in human skin. J. Dermatol. Sci. 55, 144–149.

Gilchrest, B.A. (1996). A review of skin ageing and its medical therapy. Br. J. Dermatol. 135, 867–875.

Halfter, W., Dong, S., Schurer, B., and Cole, G.J. (1998). Collagen XVIII is a basement membrane heparan sulfate proteoglycan. J. Biol. Chem. 273, 25404–25412.

Haniffa, M.A., Wang, X.-N., Holtick, U., Rae, M., Isaacs, J.D., Dickinson, A.M., Hilkens, C.M.U., and Collin, M.P. (2007). Adult Human Fibroblasts Are Potent Immunoregulatory Cells and Functionally Equivalent to Mesenchymal Stem Cells. J. Immunol. 179, 1595–1604.

Haydont, V., Neiveyans, V., Fortunel, N.O., and Asselineau, D. (2019). Transcriptome profiling of human papillary and reticular fibroblasts from adult interfollicular dermis pinpoints the “tissue skeleton” gene network as a component of skin chrono-ageing. Mech. Ageing Dev. 179, 60–77.

Janson, D.G., Saintigny, G., Van Adrichem, A., Mahé, C., and El Ghalbzouri, A. (2012). Different gene expression patterns in human papillary and reticular fibroblasts. J. Invest. Dermatol. 132, 2565–2572.

Kadler, K.E., Baldock, C., Bella, J., and Boot-Handford, R.P. (2007). Collagen at a glance. J. Cell Sci. 120, 1955–1958.

Kendall, R.T., and Feghali-Bostwick, C.A. (2014). Fibroblasts in fibrosis: novel roles and mediators. Front. Pharmacol. 5, 123.

Kolodziejczyk A.A., Kim J.K., Svensson V., Marioni J.C., Teichmann S.A. (2015). The technology and biology of single-cell RNA sequencing. Mol Cell. 58, 610–620.

Kozieł, R., Greussing, R., Maier, A.B., Declercq, L., and Jansen-Dürr, P. (2011). Functional interplay between mitochondrial and proteasome activity in skin aging. J. Invest. Dermatol. 131, 594–603.

Kretzschmar, K., and Watt, F.M. (2012). Lineage tracing. Cell 148, 33–45.

Kuroda, Y., Kitada, M., Wakao, S., Nishikawa, K., Tanimura, Y., Makinoshima, H., Goda, M., Akashi, H., Inutsuka, A., Niwa, A., et al. (2010). Unique multipotent cells in adult human mesenchymal cell populations. Proc. Natl. Acad. Sci. USA 107, 8639–8643.

Lee, S.-J., Park, C., Lee, J.Y., Kim, S., Kwon, P.J., Kim, W., Jeon, Y.H., Lee, E., and Yoon, Y. (2015). Generation of pure lymphatic endothelial cells from human pluripotent stem cells and their therapeutic effects on wound repair. Sci. Rep. 5, 11019.

Leeuwenberg, J.F., Smeets, E.F., Neefjes, J.J., Shaffer, M. a, Cinek, T., Jeunhomme, T.M., Ahern, T.J., and Buurman, W. a (1992). E-selectin and intercellular adhesion molecule-1 are released by activated human endothelial cells in vitro. Immunology 77, 543–549.

Li, A., Wei, Y., Hung, C., and Vunjak-Novakovic, G. (2018). Chondrogenic properties of collagen type XI, a component of cartilage extracellular matrix. Biomaterials 173, 47–57.

Lynch, M.D., and Watt, F.M. (2018). Fibroblast heterogeneity: implications for human disease. J. Clin. Invest. 128, 26–35.

Macosko, E.Z., Basu, A., Satija, R., Nemesh, J., Shekhar, K., Goldman, M., Tirosh, I., Bialas, A.R., Kamitaki, N., Martersteck, E.M., et al. (2015). Highly Parallel Genome-wide Expression Profiling of Individual Cells Using Nanoliter Droplets. Cell 161, 1202–1214.

Mine, S., Fortunel, N.O., Pageon, H., and Asselineau, D. (2008). Aging alters functionally human dermal papillary fibroblasts but not reticular fibroblasts: A new view of skin morphogenesis and aging. PLoS One 3, e4066.

Moll, R., Divo, M., and Langbein, L. (2008). The human keratins: biology and pathology. Histochem. Cell Biol. 129, 705–733.

Nauroy, P., Barruche, V., Marchand, L., Nindorera-Badara, S., Bordes, S., Closs, B., and Ruggiero, F. (2017). Human Dermal Fibroblast Subpopulations Display Distinct Gene Signatures Related to Cell Behaviors and Matrisome. J. Invest. Dermatol. 137, 1787–1789.

Paquet-Fifield, S., Schlüter, H., Li, A., Aitken, T., Gangatirkar, P., Blashki, D., Koelmeyer, R., Pouliot, N., Palatsides, M., Ellis, S., et al. (2009). A role for pericytes as microenvironmental regulators of human skin tissue regeneration. J. Clin. Invest. 119, 2795–2806.

Pawlina, W., and Ross, M.H. (2016). Histology: A Text and Atlas with Correlated Cell and Molecular Biology. 7th Ed. (Philadelphia: Wolters Kluwer).

Peltonen, S., Hentula, M., Hägg, P., Ylä-Outinen, H., Tuukkanen, J., Lakkakorpi, J., Rehn, M., Pihlajaniemi, T., and Peltonen, J. (1999). A novel component of epidermal cell-matrix and cell-cell contacts: Transmembrane protein type XIII collagen. J. Invest. Dermatol. 113, 635–642.

Philippeos, C., Telerman, S.B., Oulès, B., Pisco, A.O., Shaw, T.J., Elgueta, R., Lombardi, G., Driskell, R.R., Soldin, M., Lynch, M.D., et al. (2018). Spatial and Single-Cell Transcriptional Profiling Identifies Functionally Distinct Human Dermal Fibroblast Subpopulations. J. Invest. Dermatol. 138, 811–825.

R Core Team (2018). R: a Language and Environment for Statistical Computing. R Foundation for Statistical Computing, Vienna, Austria. http://www.R-project.org/

Rinkevich, Y., Walmsley, G.G., Hu, M.S., Maan, Z.N., Newman, A.M., Drukker, M., Januszyk, M., Krampitz, G.W., Gurtner, G.C., Lorenz, H.P., et al. (2015). Identification and isolation of a dermal lineage with intrinsic fibrogenic potential. Science 348, aaa2151.

Rittié, L., and Fisher, G.J. (2015). Natural and sun-induced aging of human skin. Cold Spring Harb. Perspect. Med. 5, a015370.

Rognoni, E., and Watt, F.M. (2018). Skin Cell Heterogeneity in Development, Wound Healing, and Cancer. Trends Cell Biol. 28, 709–722.

Salzer, M.C., Lafzi, A., Berenguer-Llergo, A., Youssif, C., Castellanos, A., Solanas, G., Peixoto, F.O., Stephan-Otto Attolini, C., Prats, N., Aguilera, M., et al. (2018). Identity Noise and Adipogenic Traits Characterize Dermal Fibroblast Aging. Cell 175, 1575–1590.

Satija, R., Farrell, J.A., Gennert, D., Schier, A.F., and Regev, A. (2015). Spatial reconstruction of single-cell gene expression data. Nat. Biotechnol. 33, 495–502.

Schafer, I.A., Pandy, M., Ferguson, R., and Davis, B.R. (1985). Comparative observation of fibroblasts derived from the papillary and reticular dermis of infants and adults: Growth kinetics, packing density at confluence and surface morphology. Mech. Ageing Dev. 31, 275–293.

Schindelin, J., Arganda-Carreras, I., Frise, E., Kaynig, V., Longair, M., Pietzsch, T., Preibisch, S., Rueden, C., Saalfeld, S., Schmid, B., et al. (2012). Fiji: an open-source platform for biological-image analysis. Nat. Methods 9, 676–682.

Schönherr, E., Beavan, L. a, Hausser, H., Kresse, H., and Culp, L. a (1993). Differences in decorin expression by papillary and reticular fibroblasts in vivo and in vitro. Biochem. J. 290, 893–899.

Schumacher, M., Schuster, C., Rogon, Z.M., Bauer, T., Caushaj, N., Baars, S., Szabowski, S., Bauer, C., Schorpp-Kistner, M., Hess, J., et al. (2014). Efficient keratinocyte differentiation strictly depends on JNK-induced soluble factors in fibroblasts. J. Invest. Dermatol. 134, 1332–1341.

Shain, A.H., and Bastian, B.C. (2016). From melanocytes to melanomas. Nat. Rev. Cancer 16, 345–358.

Sriram, G., Bigliardi, P.L., and Bigliardi-Qi, M. (2015). Fibroblast heterogeneity and its implications for engineering organotypic skin models in vitro. Eur. J. Cell Biol. 94, 483–512.

Simpson, C.L., Patel, D.M., and Green, K.J. (2011). Deconstructing the skin: Cytoarchitectural determinants of epidermal morphogenesis. Nat. Rev. Mol. Cell Biol. 12, 565–580.

Sorrell, J.M. (2004). Fibroblast heterogeneity: more than skin deep. J. Cell Sci. 117, 667–675.

Sorrell, J.M., and Caplan, A.I. (2009). Fibroblasts-A Diverse Population at the Center of It All. Int. Rev. Cell Mol. Biol. 276, 161–214.

Tabib, T., Morse, C., Wang, T., Chen, W., and Lafyatis, R. (2017). SFRP2/DPP4 and FMO1/LSP1 Define Major Fibroblast Populations in Human Skin. J. Invest. Dermatol. 138, 802–810.

Taniguchi, K., Arima, K., Masuoka, M., Ohta, S., Shiraishi, H., Ontsuka, K., Suzuki, S., Inamitsu, M., Yamamoto, K.I., Simmons, O., et al. (2014). Periostin controls keratinocyte proliferation and differentiation by interacting with the paracrine IL-1α/IL-6 loop. J. Invest. Dermatol. 134, 1295–1304.

Tigges, J., Krutmann, J., Fritsche, E., Haendeler, J., Schaal, H., Fischer, J.W., Kalfalah, F., Reinke, H., Reifenberger, G., Stühler, K., et al. (2014). The hallmarks of fibroblast ageing. Mech. Ageing Dev. 138, 26–44.

Veit, G., Zwolanek, D., Eckes, B., Niland, S., Käpylä, J., Zweers, M.C., Ishada-Yamamoto, A., Krieg, T., Heino, J., Eble, J.A., et al. (2011). Collagen XXIII, novel ligand for integrin α2β1in the epidermis. J. Biol. Chem. 286, 27804–27813.

Vento-Tormo, R., Efremova, M., Botting, R.A., Turco, M.Y., Vento-Tormo, M., Meyer, K.B., Park, J.-E., Stephenson, E., Polański, K., Goncalves, A., et al. (2018). Single-cell reconstruction of the early maternal–fetal interface in humans. Nature 563, 347–353.

Waldera Lupa, D.M., Kalfalah, F., Safferling, K., Boukamp, P., Poschmann, G., Volpi, E., Götz-Rösch, C., Bernerd, F., Haag, L., Huebenthal, U., et al. (2015). Characterization of Skin Aging– Associated Secreted Proteins (SAASP) Produced by Dermal Fibroblasts Isolated from Intrinsically Aged Human Skin. J. Invest. Dermatol. 135, 1954–1968.

Wang, W., Olson, D., Liang, G., Franceschi, R.T., Li, C., Wang, B., Wang, S.S., and Yang, S. (2012). Collagen XXIV (Col24a1) promotes osteoblastic differentiation and mineralization through TGF-β/smads signaling pathway. Int. J. Biol. Sci. 8, 1310–1322.

Werner, S., Krieg, T., and Smola, H. (2007). Keratinocyte-fibroblast interactions in wound healing. J. Invest. Dermatol. 127, 998–1008.

Wong, V.W., Sorkin, M., Glotzbach, J.P., Longaker, M.T., Gurtner, G.C. (2011). Surgical approaches to create murine models of human wound healing. J. Biomed. Biotechnol. 2011, 969618.

Woo, S.H., Lumpkin, E.A., and Patapoutian, A. (2015). Merkel cells and neurons keep in touch. Trends Cell Biol. 25, 74–81.

Zhang, S., and Duan, E. (2018). Fighting against Skin Aging: The Way from Bench to Bedside. Cell Transplant. 27, 729–738.

Zhuang, Y., and Lyga, J. (2014). Inflammaging in skin and other tissues - the roles of complement system and macrophage. Inflamm. Allergy Drug Targets 13, 153–161.

