## Supplementary data for "Single-cell transcriptomes of the aging human skin reveal loss of fibroblast priming"

**Figure S1**

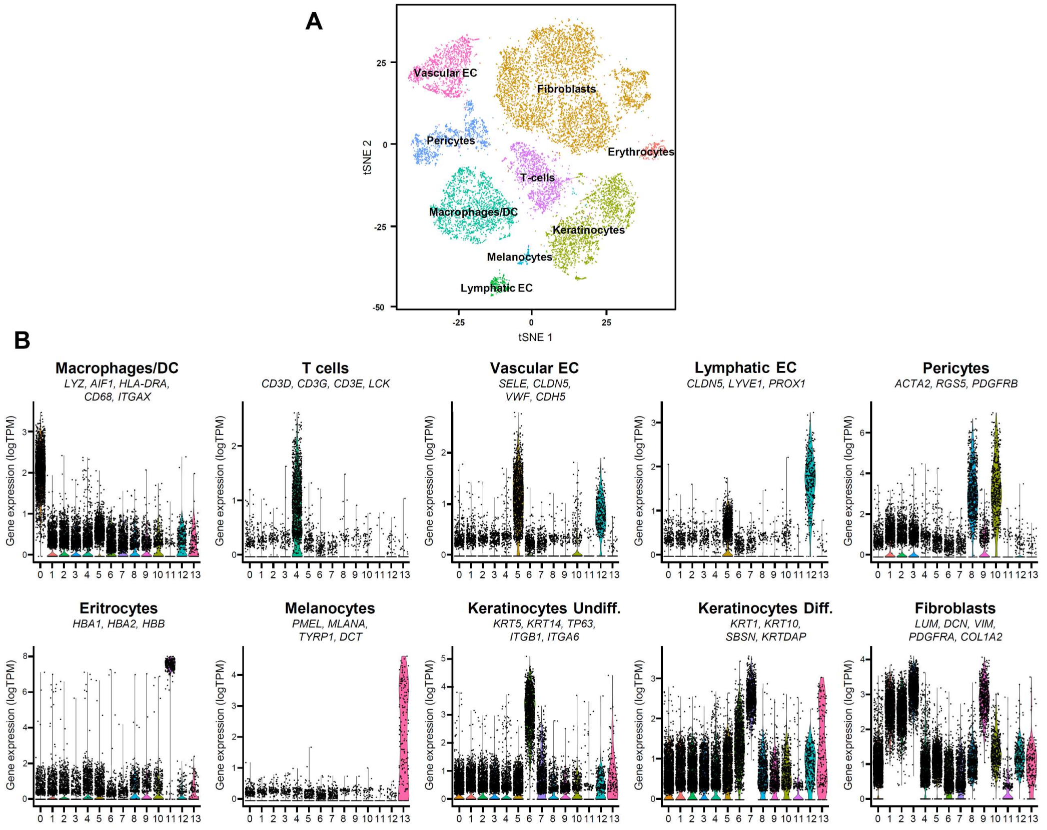


**Figure S1. Single cell RNA-sequencing of sun-protected human skin identified 9 cell types. (A)** t-SNE plot showing the distinct cell populations identified in our dataset. **(B)** Violin plots showing the average expression level of the specific markers used in Figure 1 for identifying the main cell populations of the skin. X axes depict cell cluster number and Y axes represent average expression of each set of genes in log(TPM). TPM: transcripts per kilobase million.

**Figure S2**

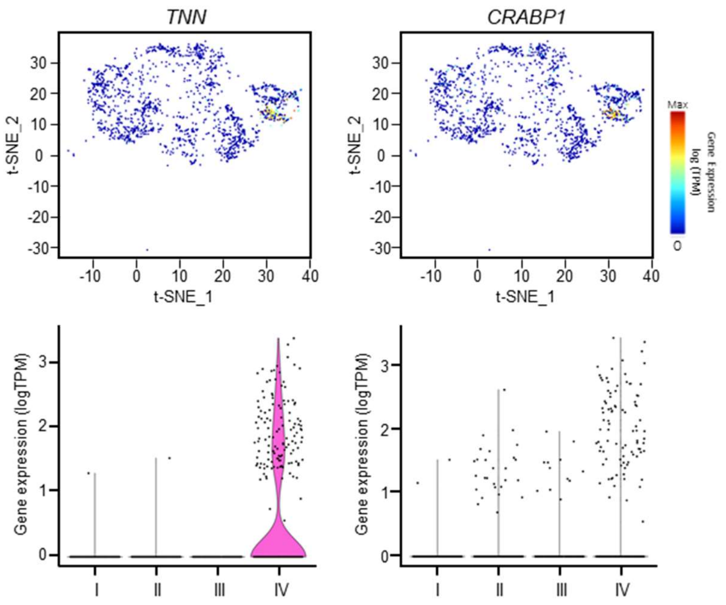


**Figure S2. Dermal Papilla (DP) stem cells are present in the mesenchymal population.** Above: gene expression of well-known DP markers *TNN* and *CRABP1* projected on the t-SNE plot of the fibroblast clusters from young skin samples (n=1,795). Red indicates maximum expression and blue depicts no expression of a particular gene. Below: Violin plots showing the expression of *TNN* and *CRABP1* in each fibroblast population in young skin. X axes depict cell cluster number and Y axes represent gene expression in log (TPM). TPM: transcripts per kilobase million.

### Figure S3

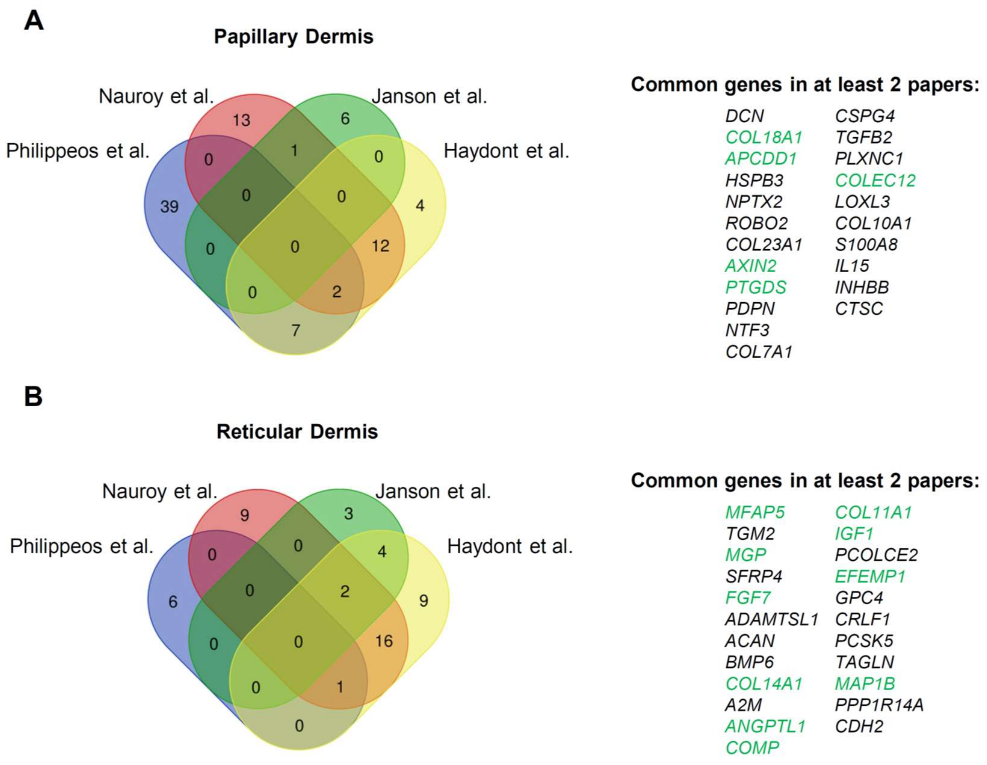


**Figure S3. Selection of gene sets defining dermal fibroblast spatial identity.** Left: Venn diagram of genes found highly expressed in the papillary **(A)** or reticular **(B)** dermis in Philippeos et al., Nauroy et al., Janson et al., and Haydont et al. Right: list of the papillary **(A)** or reticular **(B)** genes detected in at least 2 of the previous studies. In green are marked the genes found expressed by young fibroblasts and used as papillary or reticular identity in the rest of the present study.

**Figure S4**

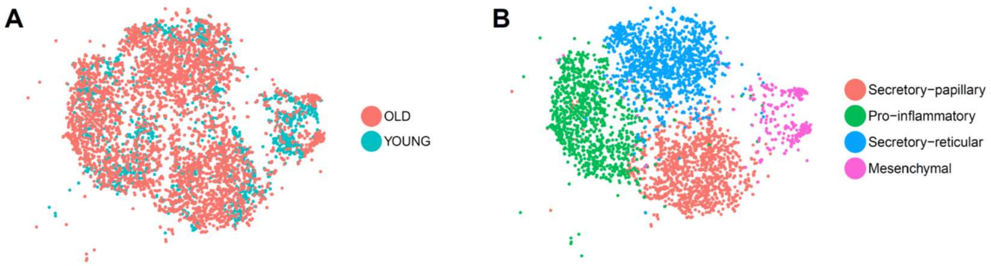


**Figure S4. Subtraction of the old fibroblasts from our complete dataset. (A)** t-SNE plot displaying dermal fibroblasts from young and old donors (n=5). Each dot represents a single cell (n=5,913). Coloring is according to age group. **(B)** t-SNE plot displaying dermal fibroblasts only from old donors (n=3). Each dot represents a single cell (n=4,118). Coloring is according to unsupervised clustering performed by Seurat.

**Figure S5**

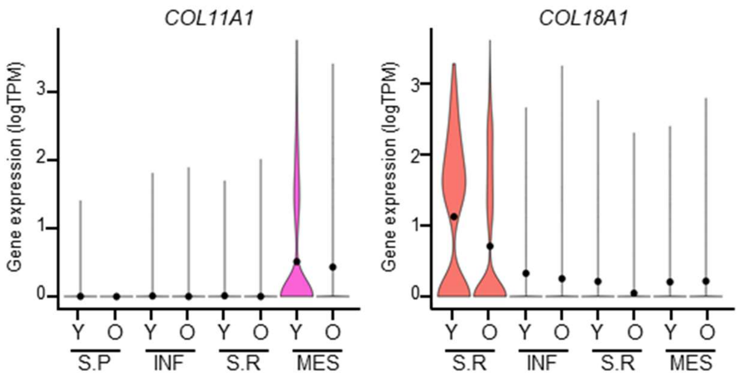


**Figure S5. Expression of functional collagens is decreased in fibroblasts upon aging.** Violin plots showing the expression of the functional collagens *COL11A1* and *COL18A1* in each fibroblast population for young and old skin. X axes depict cell cluster number and Y axes represent gene expression in log (TPM). Cluster average gene expression is shown as a black dot. TPM: transcripts per kilobase million; Y: young; O: old; S.P: Secretory-Papillary; INF: Pro-inflammatory; S.R: Secretory-Reticular; MES: Mesenchymal.

**Figure S6**

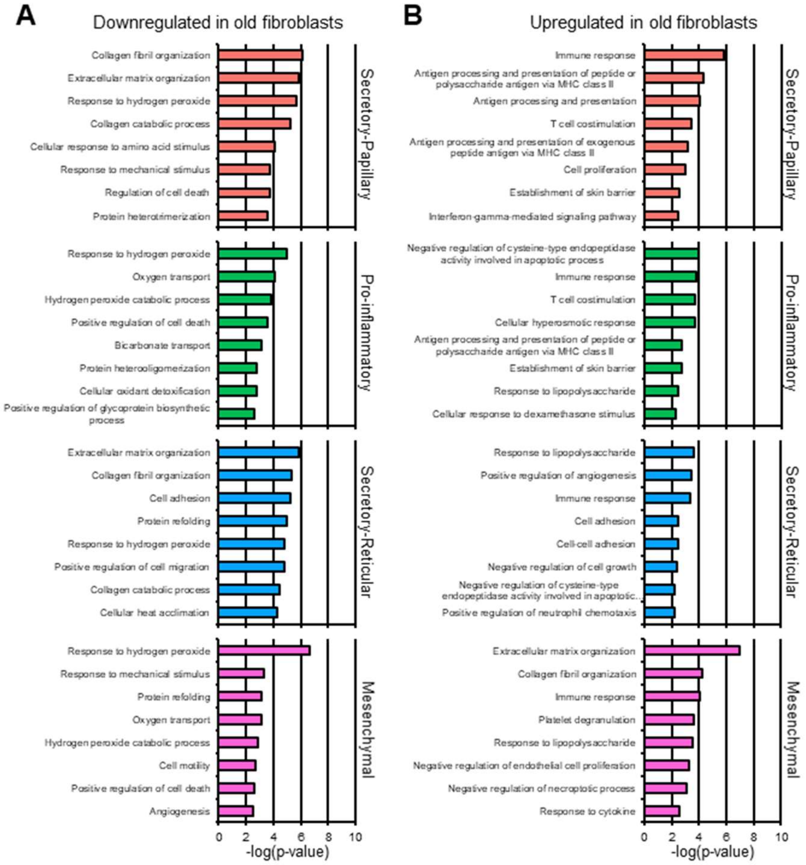


**Figure S6. Age-related changes in gene expression revel a global increase of immune response and decrease hydrogen peroxide metabolism. (A)** Top 8 enriched Gene Ontology (GO) terms obtained with the most downregulated genes in each old fibroblast population, sorted by *p*-value. **(B)** Top 8 enriched Gene Ontology (GO) terms obtained with the most upregulated genes in each old fibroblast population, sorted by *p*-value.

**Figure S7**

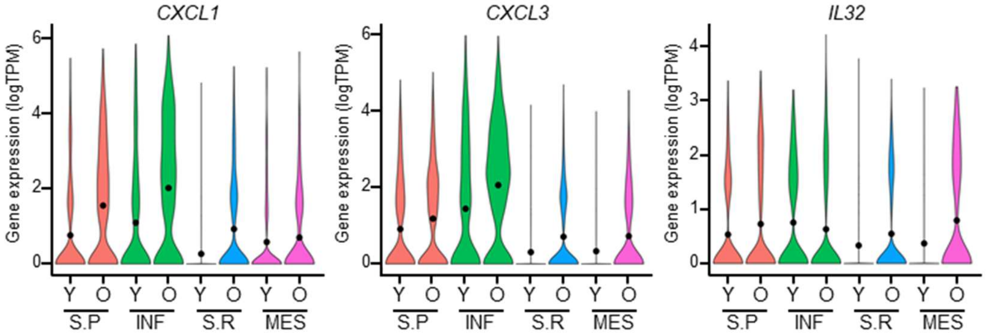


**Figure S7. Globally increased expression of pro-inflammatory cytokines in old fibroblasts.** Violin plots showing the expression of the proinflammatory cytokines *CXCL1*, *CXCL3*, and *IL32* in each fibroblast population for young and old skin. X axes depict cell cluster number and Y axes represent gene expression in log (TPM). Cluster average gene expression is shown as a black dot. TPM: transcripts per kilobase million; Y: young; O: old; S.P: Secretory-Papillary; INF: Pro-inflammatory; S.R: Secretory-Reticular; MES: Mesenchymal.

**Figure S8**

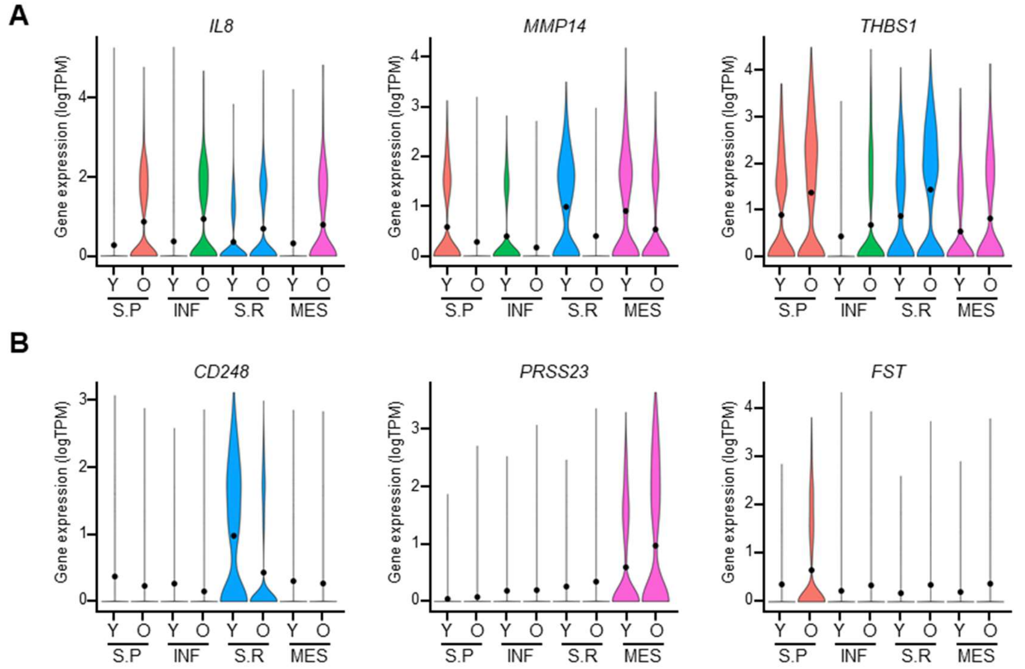


**Figure S8. Old fibroblast populations express distinct SAASP-associated genes. (A)** SAASP-associated genes that change their expression upon aging in the same manner in all fibroblasts populations. Violin plots show the expression of *IL8*, *MMP14* and *THBS1* in each fibroblast population for young and old skin. **(B)** SAASP-associated genes that change their expression upon aging specifically in only one fibroblast population. Violin plots show the expression of *CD248*, *PRSS23* and *FST* in each fibroblast population for young and old skin. X axes depict cell cluster number and Y axes represent gene expression in log (TPM). Cluster average gene expression is shown as a black dot. TPM: transcripts per kilobase million; Y: young; O: old; S.P: Secretory-Papillary; INF: Pro-inflammatory; S.R: Secretory-Reticular; MES: Mesenchymal.

**Figure S9**

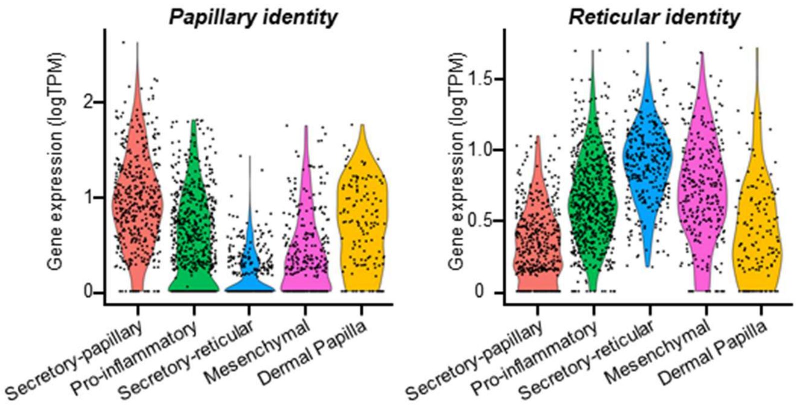


**Figure S9. Dermal papilla stem cells show an enriched papillary identity.** Average expression of the genes constituting the papillary and reticular gene signatures for predicting dermal localization of the fibroblast subpopulations. Dermal papilla stem cells were subtracted from the mesenchymal group. X-axes depict cell cluster number and Y-axes represent gene expression in log(TPM). TPM: transcripts per kilobase million.

**Figure S10**

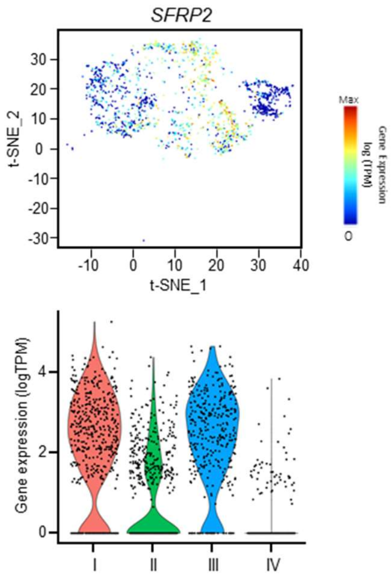


**Figure S10. Expression of *SFRP2* is enriched in the secretory populations.** t-SNE and violin plots showing the expression levels of *SFRP2* in the fibroblast subpopulations. In the t-SNE plots, red indicates maximum relative expression and blue indicates low or no expression of a particular gene. In the violin plot, the X-axis depicts cell cluster identity while the Y-axis represents gene expression in log(TPM). TPM: transcripts per kilobase million.

| Sample ID | Gender | Age | Skin type | Reads per sample | N° of cells | Reads per cell | Genes per cell |
|---|---|---|---|---|---|---|---|
| Young 1 | Male | 25 | Fair | 322,091,192 | 2784 | 102,904 | 1343 |
| Young 2 | Male | 27 | Fair | 338,738,780 | 2670 | 119,737 | 1111 |
| Old 1 | Male | 53 | Fair | 359,776,321 | 3324 | 107,976 | 1718 |
| Old 2 | Male | 70 | Fair | 378,219,220 | 2144 | 170,215 | 1388 |
| Old 3 | Male | 69 | Fair | 370,342,531 | 4535 | 81,411 | 872 |

**Table S1. Summary of the samples included in this study and their main characteristics.** Table depicts biological features (gender, age at the time of excision and type of skin) and sequencing technicalities (reads/sample, nº of cells, reads/cells and genes/cell) of our five samples which were stratified according to age (sample ID).
